# Single-cell transcriptional profiling reveals cellular and molecular divergence in human maternal-fetal interface

**DOI:** 10.1101/2022.02.15.479798

**Authors:** Quanlei Wang, Jinlu Li, Shengpeng Wang, Qiuting Deng, Yanru An, Yanan Xing, Xi Dai, Zelong Li, Qiwang Ma, Kuixing Wang, Chuanyu Liu, Yue Yuan, Guoyi Dong, Tao Zhang, Huanming Yang, Yutao Du, Yong Hou, Weilin Ke, Zhouchun Shang

**Author notes:** Correspondence and requests for materials should be addressed to W.K. or to Z.S. These authors contributed equally to this paper (Quanlei Wang, Jinlu Li, Shengpeng Wang).

## Abstract

Placenta play essential role in successful pregnancy, as the most important organ connecting and interplaying between mother and fetus. However, the cellular and molecular characteristics of fetal origin and maternal origin cell populations within the fetomaternal interface still is poorly understood. Here, we profiled the transcriptomes of single cells with well-defined maternal-fetal origin that consecutively localized from fetal section (FS), middle section (Mid_S) to maternal section (Mat_S) within the human full-term placenta. Then, we initially identified the cellular and molecular heterogeneity of cytotrophoblast cell (CTB) and stromal cell (STR) with the spatial location and fetal/maternal origin, also highlighted STR cells from fetal origins showed greater proliferation ability in Mat_S compared to cells from FS or Mid_S. Further, by integrating analysis with the first-trimester placental single cell transcriptome data, we revealed that a subpopulation of trophoblast progenitor-like cells (TPLCs) existed in the full-term placenta and mainly distributed in Mid_S, with high expression of pool of putative cell surface makers and unique molecular features. Moreover, through the extravillous cytotrophoblast (EVT) subsets differentiation trajectory and regulation network analysis, we proposed a putative key transcription factor *PRDM6* that promoted the differentiation of endovascular extravillous trophoblast cells (enEVT). Finally, based on the integrated analyses of single cell transcriptional profiling of preeclampsia (PE) and match-trimester normal placenta, we highlighted the defective EVT subgroup composition and down-regulation of *PRDM6* may lead to an abnormal enEVT differentiation process in PE. Together, our study offers important resources for better understanding of human placenta, stem cell-based therapy as well as PE, and provides new insights on the study of tissue heterogeneity, the clinical prevention and control of PE as well as the maternal-fetal interface.

## Introduction

Human placenta is a complex anatomic structure derived from trophectoderm and extraembryonic mesoderm^1^. It is responsible for regulating immune system and transporting nutrients and waste between fetus and mother. Various specialized cells derived from fetal and maternal with coordinated mRNA transcriptional regulation during human placentation and maturation contribute to this vital task^1, 2^. Any cellular and molecular abnormality in the maternal-fetal interface may lead to multiple pregnancy outcomes, such as preeclampsia (PE), which are leading causes of maternal and neonatal death^3–5^. The maternal-fetal interface is generally consecutive from fetal side to maternal side with corresponding fetal or maternal origin cell types distribution^1^. For instance, some fetal derived trophoblast cells mainly located in fetal side, also migrated to maternal side for placental anchoring and tissue remodeling. On the other hand, previous study reported that the fetal side also infiltrate maternal derived cells, including placenta chorionic villus, chorionic plate and chorionic membrane through the intervillous space^6^. For other cell types, abundantly resided in maternal-fetal interface, e.g., stromal cells (STR) from both fetal and maternal origin, play crucial roles in modulating multicellular interaction by releasing signal molecules. And, STR culture-expansion *in vitro* holds great promising in regenerative medicine. Current, human placenta has been regarded as an ideal tissue source for STR isolation and preservation in biobank^7, 8^. However, the molecular features and functional differences of primary STR with specific origin and spatial location in the maternal-fetal interface still remain unclear.

Based on current knowledge, the trophoblast cells from placenta include three major functional cell populations: cytotrophoblast (CTB), syncytiotrophoblast (STB) and extravillous trophoblast (EVT). Previous studies showed the proliferative CTB as the initial cell population for STB and EVT differentiation during early placenta development. Large studies showed trophoblast progenitor cells (TPCs) existed in placenta early villus CTB, but rapidly decreased after first-trimester stage^9–11^. Also, several studies have successfully isolated TPCs as cell culture model from the first-trimester placenta villus tissue^12^ or from the differentiation of pluripotent stem cells *in vitro*^13, 14^. However, whether TPCs exist in human full-term placenta is still undetermined.

Out of chorionic villus, the EVT populations that originated from CTB, undergoing serially differentiation and migration to remodel endometrium and spiral artery in maternal tissue to ensure blood flow circulation. The EVT differentiation from CTB is a complex process, and including multiple subpopulations that responsible for specific functional fate. Based on current knowledge, the proliferation CTB form extravillous trophoblast cells column (column EVT) at the tip of villus, then, the column EVT differentiate further into interstitial extravillous trophoblast cells (iEVT) and endovascular extravillous trophoblast cells (enEVT) for invading endometrium and spiral artery respectively. At present, existed several markers are used to distinguish the EVT subpopulations described above, such as *MKI67* for column EVT, *ITGA1* for iEVT and enEVT. However, we still know little about transcriptional regulation and pathways involved in EVT differentiation and invasion, especially, the regulation of both iEVT and enEVT under normal condition and pregnancy-related diseases.

Single-cell RNA-sequencing (scRNA-seq) technologies have greatly improved our understanding of heterogeneity in terms of cell fate determination and transcriptional regulation of development^15–17^. Current, several studies have performed human maternal-fetal interface single cell transcriptome analysis, but most of them focused on the first-trimester pregnancies or integrated analysis of cell lineages without specific origin and spatial location^18–20^. For instance, Roser Vento-tormo et al. revealed the cellular heterogeneity of the first-trimester placenta, and develop a repository of ligand–receptor complexes that are critical for placentation and reproductive success^20^. Moreover, Pavličev *et al*. inferred the cell-cell interactome by assessing the gene expression of ligand-receptor pairs across cell types and found that highly cell-type specific expression of a group of G-protein-coupled receptors could be a reliable tool for cell type identification from 87 single-cell transcriptomes. They also suggested that uterine decidual cells represent a cell-cell interaction hub with a large number of potential signal exchange. Growth factors and immune signals dominate among the transmitted signals, which suggest a delicate balance of enhancing and suppressive signals^21^. Tsang *et al*. dissected the cellular heterogeneity of the human placenta and defined individual cell-type specific gene signatures by analyzing nonmarker selected placenta cells from third-trimester placenta and preeclamptic placentas using large-scale microfluidic single-cell transcriptomic technology^22^. Overall, previous studies showed accurate cellular atlas for early stage of human placenta development, but that for the full-term placenta is largely lacking. Moreover, both the regulatory mechanism of trophoblast subpopulations differentiation and interactions between cell types within the maternal-fetal interface still remains elusive.

In the present study, we profiled the transcriptomes of single cells that consecutively localized from fetal section (FS), middle section (Mid_S) and maternal section (Mat_S) of human full-term placenta based on previous study^1^. We dissected cell populations with indication of their fetal or maternal origin base on single-cell SNV analysis. Then, we observed the spatial variation of cellular composition from the FS, Mid_S to Mat_S, and highlighted the molecular and functional diversities of CTB and STR. Moreover, we integrated the first-trimester placental single cell transcriptome data with our trophoblast cells and reconstructed the differentiation relationships within the trophoblast subtypes, then revealed trophoblast progenitor-like cells (TPLCs) with unique molecular feature mainly distributed in the Mid_S. Additionally, we proposed putative key transcription factors, *PRDM6* (PR/SET domain 6) that may play critical role in promoting enEVT differentiation through cell-cycle arrest signals. Finally, compared with the transcriptional profiling of the normal placenta tissues, the PE placenta showed abnormal epithelial-to-mesenchymal transition related ligand-receptor interactions and down-regulation of *PRDM6* may lead to dysregulated enEVT differentiation. Collectively, these results not only offer insights into the spatial structure and function of human placenta but also provide an important resource that will pave the way for basic research and regenerative medicine in placental development field.

## Results

### Dissecting maternal and fetal cell heterogeneity in human full-term fetal-maternal interface

Total 11,438 droplet-based single cell transcriptomes of human full-term placenta were harvested with consecutive spatial locations, including fetal section (FS), middle section (Mid_S) and maternal section (Mat_S) (Fig. 1a, Supplementary Fig. 1a). Unsupervised graph-based clustering of the dataset was performed to produce 27 clusters after computational quality control (see Methods). Cluster-specific expression pattern of known marker genes was employed to annotate the major cell types including villous cytotrophoblasts (CTB; marked by *KRT7*, *PAGE4*, *GATA3*), extravillous trophoblasts (EVT; *HLA-G*, *PAPPA2*), syncytiotrophoblasts (STB; *CGA*, *CYP19A1*) , stromal cells (STR; *THY1*, *DCN*) , decidual cells (DEC; *DKK1*, *IGFBP1*), perivascular cells (PV; *MYH11*, *NDUFA4L2*), vascular endothelial cells (VEC; *PECAM1*, *IFI27*) , lymphatic endothelial cells (LEC; *LYVE1*, *CCl5*), and immune cells (IMM; *PTPRC*, *CD74*) (Fig. 1b, 1c, 1d; Supplementary Fig. 1b, 1c, 1d). These cells showed significant cellular heterogeneity which was consistent with previous bulk RNA sequencing data and single cell transcriptomic profiling of biopsies taken from different areas of the placenta interface^1, 18^.

**Fig. 1.**
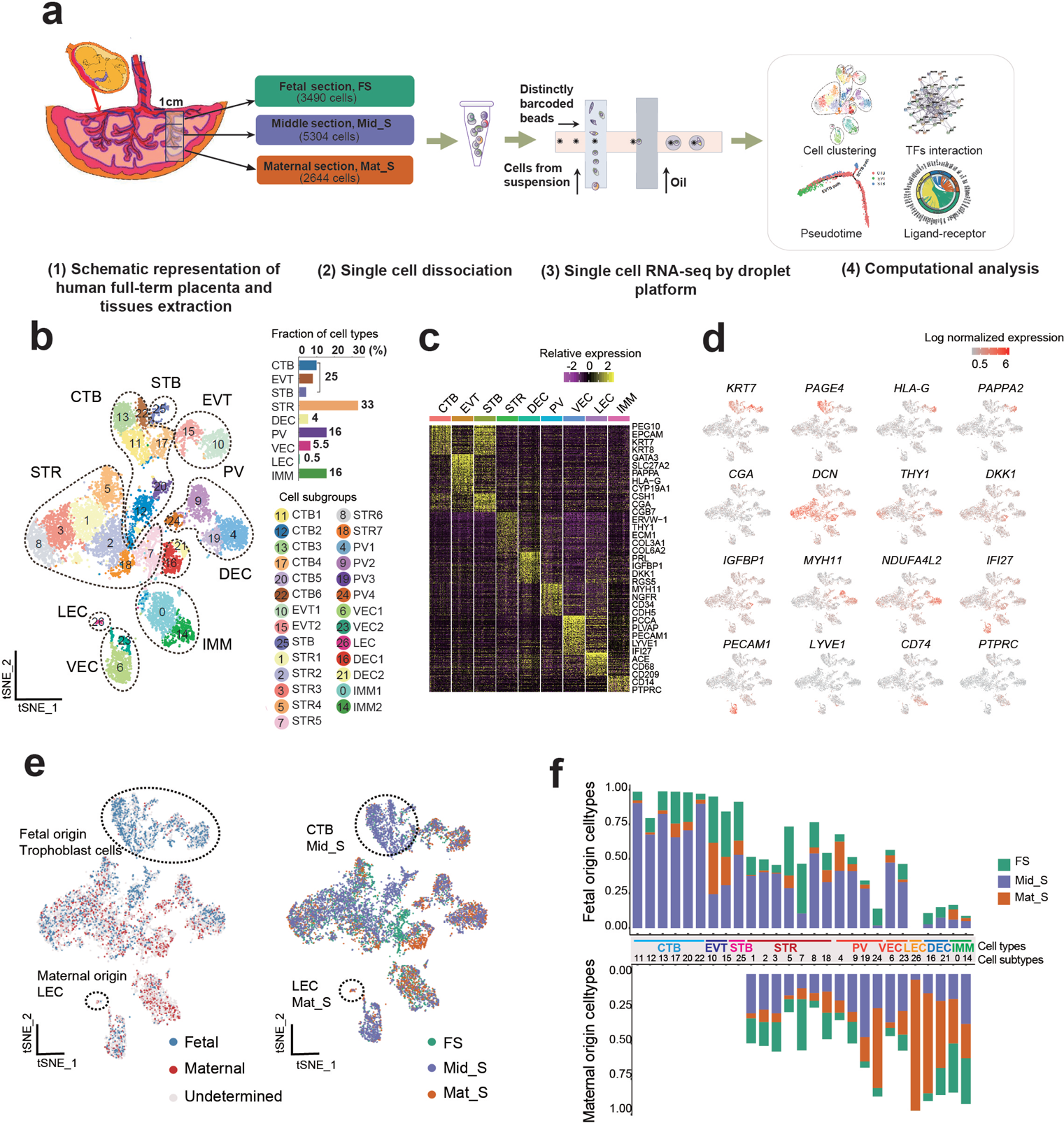
Dissecting cellular heterogeneity of human full-term placenta. a. Workflow of single-cell transcriptome profiling of human full-term placenta. b. t-SNE analysis of human full-term placenta (Left). Each dot represents an individual cell. Colors indicate cell type or state. PV, perivascular cell; STR, stromal cell; IMM, immune cell; CTB, villous cytotrophoblast; EVT, extravillous trophoblast; STB, syncytiotrophoblast; VEC, vascular endothelial cell; LEC, lymphatic endothlial cell; DEC, decidual cell. The column chart shows the fraction of indicated cell types (Right). c. Heatmap shows the top differentially expressed genes of each cell type. Color scheme is based on relative gene expression (z-score). d. t-SNE plot showing the selected cell type-specific marker gene expression pattern in human placenta. e. Origin (Left) and location (Right) of each cell are shown using the same layout as in figure 1b. Circle mark cell types with relatively specific origin or spatial localization. f. Column chart shows the percentage of indicated cell types from fetal or maternal origin in specific spatial location, respectively.

To further distinguish the maternal or fetal origin of single cells within the full-term placenta using previous reported method^20^. The ratio of Mahalanobis distance of fetal cells, maternal cells and assigned cells of fetal or maternal origin were calculated accordingly using the difference ratio between a single cell SNV and the corresponding fetal SNV datasets reference (Fig. 1e, 1f; Supplementary Fig. 1e; see Methods). The results show that maternal cells including LEC and DEC mainly dominated the Mat_S; CTB, EVT, and STB were derived from fetal origin and mainly distributed in Mid_S; proportionate STR, PV, and VEC originated from both fetal and maternal compartments which mainly occupied Mid_S; IMM mixed with fetal and maternal origin distributed in each section proportionally. The fetal and maternal orgin identifies was similar with that in first-trimester placenta in previous study^20^. In additional, a more comprehensive cellular map with fetal and maternal orgin and spatial distribution of the full-term fetal-maternal interface was established in our study.

### CTB and STR molecular and functional diversity within spatial location and origin

To further dissect the cellular heterogeneity of specific spatial location within placenta interface. Cells from FS, Mid_S and Mat_S areas were re-clustered while each cluster was annotated with well-known cell type markers respectively. As expected, multiple CTB subpopulations were identified within each spatial section (Fig. 2a). Among these CTB subpopulation, one subpopulation in the Mid_S is high expression of cell-cycle related gene *MKI67*, suggesting that highly proliferative CTB also exist in specific location of full-term placenta (Fig. 2b). Then, GO term enrichment analysis was performed for CTB in FS, Mid_S, and Mat_S, respectively. As expected, these GO terms generally divided into common and spatial section-specific groups, for the common terms included “placenta development”, “female pregnancy”, “embryo implantation”, and “post-embryonic animal morphogenesis”, which indicated that the fundamental functions of the placenta were revealed by our data analysis. Then, for the spatial section-specific group terms, for instance, CTB in FS, as the outermost part of placenta and side of umbilical cord insertion, enriched GO terms like “cellular response to gamma radiation”, “regulation of oxidative phosphorylation”, and “cellular response to X-ray”, with high expression of *EGR1* and *TGFBI,* which involve in regulating radiation-induced cell activity have been reported, previously^23–25^. Also, CTB in Mid_S showed high expression of *PRDX2* and *SPINT2* with “positive regulation of exosomal secretion”, “extracellular vesicle biogenesis”, “extracellular exosome biogenesis”, and “exosomal secretion” were enriched (Fig. 2c). Above terms are expected as Mid_S, the location for metabolites exchange between fetus and mother and in line with previous studies^26, 27^. In addition, CTB in Mat_S showed high expression of *MAP2K3* and *XBP1* while the enriched terms include “positive regulation of inflammatory response”, “endothelial cell migration”, and “regulation of vasculature development” (Fig. 2c, 2d, 2e). Based on the above findings, we infer that the human placenta performs executive function through specific trophoblast cell population, and here, our study indicated that CTB populations perform multiple functions via specific spatial microenvironment with specific molecular enrichment expression in the interface. Collectively, we provided a precise study of cellular molecular features of CTB subpopulation with structure and spatial location in the interface, and opening a window with higher resolution for deeper understanding of trophoblast subpopulation biological activities and fetal development.

**Fig. 2.**
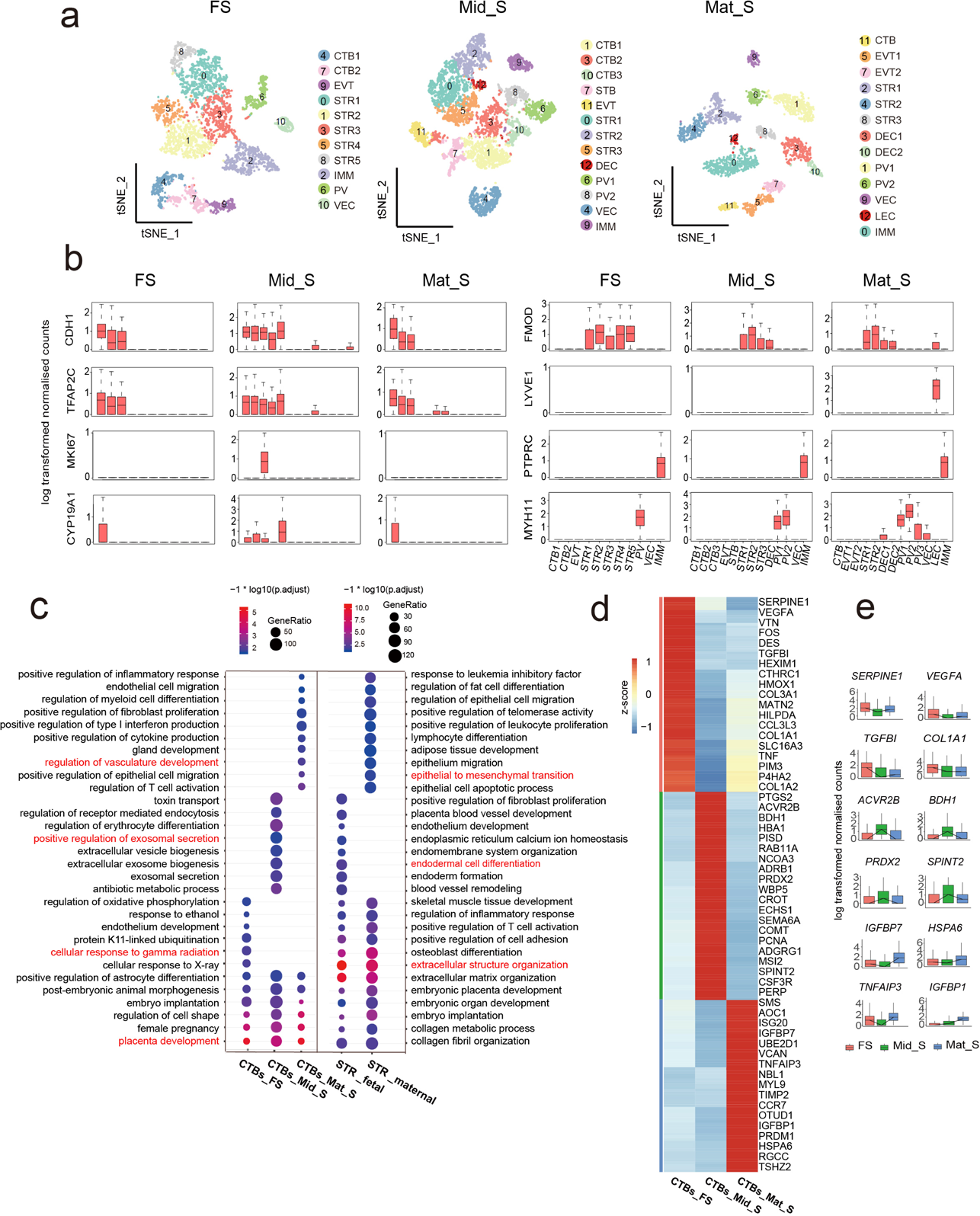
Reconstruction of spatial heterogeneity of cell type and gene expression pattern in the maternal-fetal interface. a. t-SNE plots shows single-cell transcriptomic clustering of three specific tissue locations (including FS, Mid_S, Mat_S) in full-term maternal-fetal interface, respectively. Each dot represents an individual cell. Cells are colored by cell-type cluster. b. Boxplot showing the relative expression levels of selected markers for each cell cluster. c. Selected GO terms identified by highly expressed genes of CTB in FS, Mid_S, and Mat_S, and STR of fetal and maternal origin, respectively. (Top 1000 highly expressed genes were selected for GO analysis. Highly expressed gene: expressed cell number > 20% and gene coefficient of variability (CV) <1 in each section). d. Heatmap showing the selected differentially expressed genes of CTB subpopulations derived from FS, Mid_S and Mat_S. Red corresponds to a high expression level; blue and black correspond to low expression level (the differentially expressed genes were identified by FindAllMarker function in Seurat, p_val_adj < 0.05; avg_logFC > 0.25) e. Boxplots showing the expression of selected genes from figure 2d.

The STR in human placenta had showed heterogeneous populations with specific spatial location and origin using traditional methodology^28^. To further address this item at single-cell resolution, GO enrichment analysis showed that STR in both fetal and maternal origin not only exhibited high biological activity involved in “extracellular matrix organization” and “collagen fibril organization”, but also showed key roles in “embryo implantation” and “embryonic organ development” (Fig. 2c). The results potentially indicated STR has crucial role in regulating placenta and embryonic development and was in line with previous studies^29, 30^. Moreover, fetal STR might be advantage in endomembrane related system development, while maternal origin STR showed great value in regulation of immune response related activity in our study (Fig. 2c). For the spatial location analysis of STR with inferred origin, to our surprise, STR both fetal and maternal STR derived from Mat_S showed higher proliferative activity through regulation of cell cycle G2/M phase transition pathway and telomere maintenance related pathway, respectively based on the GO enrichment analysis (Supplementary Fig. 2a). Also, the stemness-related genes including *THY1* and *VCAM1* were highly expressed in Mat_S STR, as well as cytokines and hormones-related genes like *PGF*, *FGF2*, *FGF10 etc*. that have crucial role in maintaining STR self-renewal and functional actives (Supplementary Fig. 2b, 2d). Furthermore, cell surface markers including *THY1*, *CD151*, *CD99*, *IL6ST*, *PDGFRA, etc*., involved in promoting STR proliferation, also helped to distinguish fetal and maternal STR in Mat_S (Supplementary Fig. 2e). These genes in line with the functional terms related to positive regulation of cell cycle of fetal STR in Mat_S, such as well-known stemness related gene such as *THY1*, *CD151* ^28, 31^, relatively higher expression in fetal STR than that in maternal STR in Mat_S (Supplementary Fig. 2a, 2e). Comparison of STR populations within the same Mat_S of placenta confirmed fetal origin likely to be more stemness than maternal origin STR. Here, for the first time we presented the whole genome wide molecular profiling differences of fetal and maternal STR in the same tissue origin, and the identified gene profiles were employed to further isolate STR with specific origin from Mat_S of full-term placenta tissue *in vitro*.

#### Trophoblast development trajectory reveals TPLCs in full-term placenta

To investigate the regulation process of trophoblast differentiation and highlight the stemness feature of trophoblasts, our single cell transcriptome data was integrated with published first-trimester placenta transcriptome data^20^. The trophoblast populations were sub-clustered into CTB subpopulations, EVT subpopulations, and STB subpopulations (Fig. 3a; Supplementary Fig. 3a, 3b, 3c). Then, the differentiation trajectory was constructed using the inferred subgroups. As expected, trophoblast cells formed a continuous “Y-shaped” trajectory, in which CTB was located at the trunk with high expression of proliferation and stemness related genes, for instance *TEAD4*, *KRT8*, and the two branch arms were occupied by the differentiation to EVT direction and STB direction (Fig. 3b, 3c; Supplementary Fig. 3d). Genes related to migration and invasion were highly expressed in the cells on EVT path, such as *HLA-G*, *PLAC8, ASCL2*, *EBI3*, *PAPPA*, and *PAPPA2*, which was consistent with previous studies^22^, while genes related to hormone and cell fusion, such as *CGA*, *ERVFRD-1*, *ERVW-1*, *LGALS16*, and *CYP19A1*, expressed in the cells on STB pathway (Fig. 3c, 3d).

**Fig. 3.**
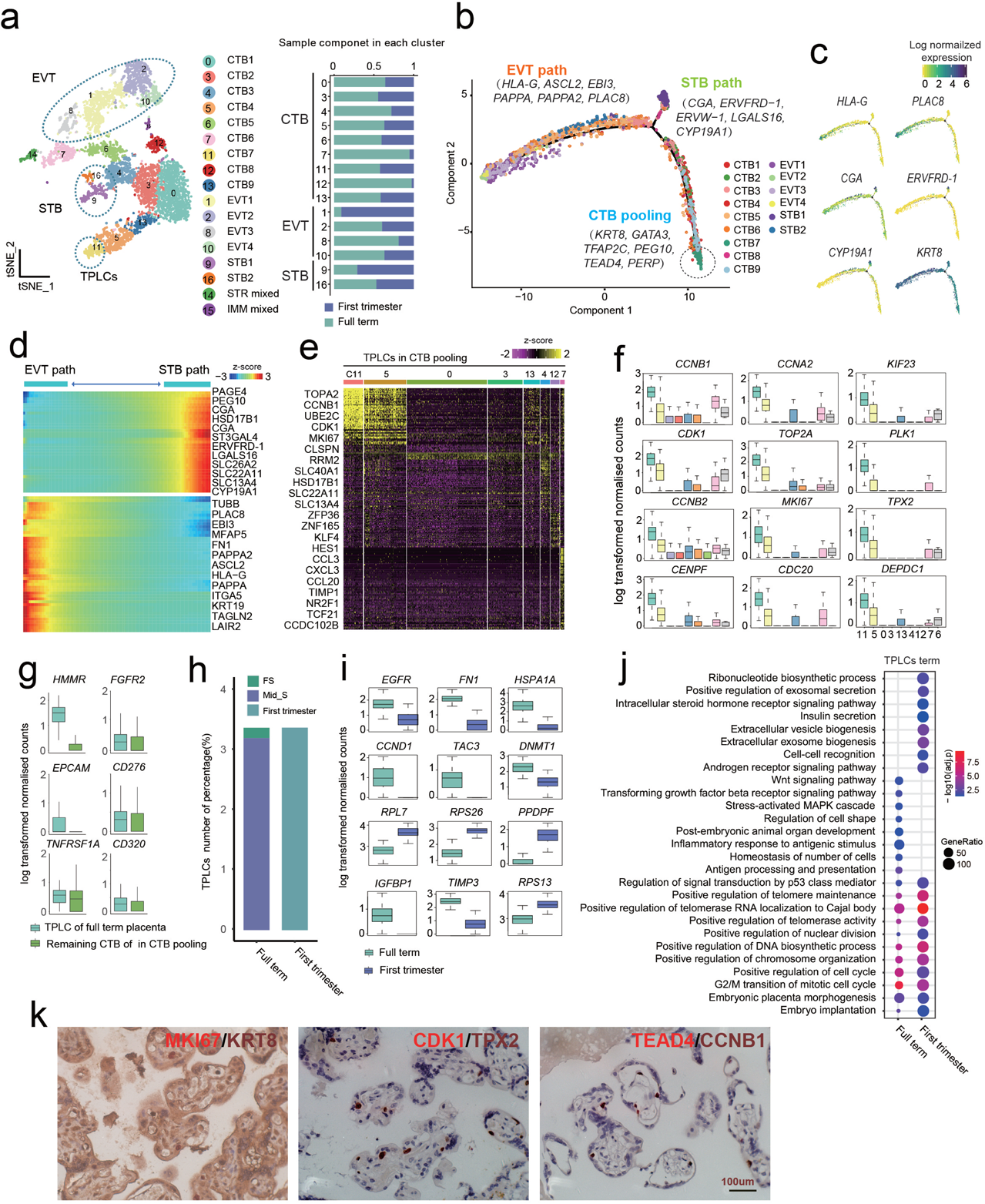
The trophoblast progenitor like cells (TPLCs) existed in human full-term placenta. a. t-SNE visualization of trophoblast cells from integrated data of full-term placenta cells and the published first-trimester placenta cells shown in Supplementary figure 3a. On the right, the barplot shows the proportion of full-term placenta cells and first-trimester placenta cells in each cluster and each cell type. b. Pseudotime ordering of trophoblast subgroups that reveals EVT and STB pathway and visualization in biaxial scatter plot. c. Expression pattern of selected genes across trophoblast differentiation branches on the reconstructed trajectory. Color scheme is based on log-transformed, normalized expression levels . d. Heatmap showing the selected differentially expressed genes expression of genes that are identified as significantly involved in EVT and STB differentiation pathway. Color scheme is based on relative gene expression (z-score). e. Heatmap shows the differentially expressed genes among CTB subpopulations, in which one small cluster (C11, termed as TPLCs) shows highly expressed cell cycle-related genes. f. Boxplot showing the log-transformed, normalized expression of genes selected from figure. 3e. g. Boxplot showing the expression level of selected cell surface genes between TPLCs and other CTB clusters derived from full-term placenta in CTB branch of figure. 3b. Two-sided Wilcoxon rank sum test were calculated, *: p < 0.05; **: p <0.01; ****: p < 0.0001 h. Column chart showing the percentage of stemness trophoblast cells derived from indicated gestation and spatial location. i. Boxplot showing the differentially expressed genes of stemness trophoblast cells derived from first-trimester and full-term placenta. Genes were selected from top 50 differentially expressed genes identified by FindAllMarker function in Seurat, p_val_adj < 0.05; avg_logFC > 0) j. GO enrichment analysis showing the selected functional terms of TPLCs derived from first-trimester and full-term placenta. k. Immunostaining of MKI67, KRT8, CDK1, TPX2, TEAD4 and CCNB1 in Mid_S of human full-term placenta. Scale bar represents 100 μm.

Interestingly, a minor subpopulation cluster11 (C11) derived from both first- and third-trimester placenta at the head of trunk on trophoblast trajectory in the CTB subpopulations also mentioned above showed highly expression of proliferative activity-related genes, e.g., *MKI67, CCNB1, CDK1* and *TOP2A,* also stemness related genes, e.g., *TEAD4*, *TPX2, TFAP2C,* suggesting the possible existence of stemness-trophoblast cells, here named: trophoblast progenitor-like cells (TPLCs) in human full-term placenta, and trophoblast progenitor cells (TPCs) in human first-trimester placenta in the present study, respectively (Fig. 3e, 3f). To further characterize TPLCs of full-term placenta in our study, we extracted the C11 cells derived from third-trimester placenta and identified the differentially expressed genes (DEGs) of each CTB subgroup and compared the gene expressions between C11 and all other CTB subgroups of full-term placenta. The results showed unique expression pattern of cell cycle-related genes in TPLCs (C11) with highly expression of cell surface maker *HMMR*. Besides, TPLCs mainly localized in the Mid_S. Furthermore, we found highly expressed genes including *EGFR, FN1, HSPA1A* and *CCND1* in TPLCs derived from full-term placentas, while *RPL7, RPS26,* and *PPDPF* were highly expressed in TPCs from first-trimester placentas. The GO enrichment analysis showed that TPCs of first-trimester maintained self-renewal and differentiation potency by two pathways, “intracellular steroid hormone receptor signaling” and “androgen receptor signaling pathway”, which play crucial roles in stem cell division and differentiation during early human embryogenesis^32^, while “Wnt signaling pathway” and “transforming growth factor beta receptor signaling pathway” were enriched in TPLCs of full term(Fig. 3i, 3j), previous reports indicated that Wnt activation and TGF-β inhibition play essential roles in long-term culture of human villous CTB^10^. Also, the immumohistochemical staining Mid_S of human placenta tissue showed that KRT8 was co-expressed with MKI67, and CDK1 was co-expressed with TPX2, and TEAD4 was co-expressed with CCNB1 in some specific cells which was consistent with the mRNA expression level (Fig. 3k). Based on above results, we proposed that some trophoblast cells simultaneously expressed proliferative and stemness related markers, might act as the TPLCs in human full-term placenta. To our knowledge, this is the first insights on TPLCs of full-term placenta, and provided gene markers based on bioinformatics analysis for isolating TPLCs from placenta as well as potential cell models application for disease mechanism research.

### Identifying key transcription factors (TFs) of EVT subpopulation differentiation and invasion

Based on trophoblast subclustering analysis, total four subclusters of HLA-G^+^ EVT were identified, including C1, C2, C8 and C10. C1 was defined as column EVT with high expression of *MKI67*, *TET1*, and *CDK1*; C2 and C8 was defined as iEVT1 with high expression of *ITGA1*, *MCAM*, and *TAC3* that related to invasion, migration, and stromal cell characteristics and iEVT2 with high expression epithelial and smooth muscle cell-related markers *PAEP*, *ACTA2*, and *TAGLN*; C10 was more likely enEVT by expressing higher levels of *ITGB1*, *CDH1*, and *CD44* that are related to extracellular structure organization compared to iEVT1(Supplementary Fig. 4a). Furthermore, the GO enrichment analysis of column EVT, iEVT1, iEVT2 and enEVT were performed by cluster-specific gene (Supplementary Fig. 4b). As expect, the terms “regulation of body fluid levels”, and “cellular response to amino acid stimulus” for iEVT2, suggest iEVT2 invading toward glands^33^. Whereas, the enEVT and iEVT1 were commonly enriched terms “extracellular structure organization” and “response to hypoxia”, by contrast, the enEVT were enriched the terms “positive regulation of blood vessel endothelial cell migration”. All the above results suggest the unique characteristics of molecular and functional state in the four EVT subclusters were observed at single cell transcriptome level of full-term placenta.

Previous studies showed that transcription factors (TFs) played crucial roles in regulating development and function of trophoblasts^34^. Understanding the TFs regulation network that guiding differentiation and invasion of EVT subgroups during placenta development is a major challenge. To investigate the regulation dynamics of TFs, we first inferred trajectories of EVT using partition-based approximate graph abstraction (PAGA) analysis based on the four EVT subpopulations mentioned above. The results showed that the column EVT localized at the starting point of trajectory and differentiated towards three directions: column EVT to iEVT1 to enEVT, column EVT to iEVT2, and column EVT to enEVT (Fig. 4a). The TFs/genes dynamically modulated within each direction of differentiation were presented (Fig. 4a). As a crucial role of enEVT in remodeling of the uterine spiral arteries, we focused on the column EVT to iEVT1 to enEVT direction and exacted two pool of TFs/genes with different expression pattern (Fig. 4b); the cell proliferation related gene, including *MKI67*, *CDK1*, *HDAC1* etc. were greatly down-regulated; while *THBS1*, *CXCL8* and *IL6* etc. expression was largely activated during enEVT differentiation (Fig. 4c). In line with the dynamical gene expression, the GO enrichment analysis demonstrated that positive regulation of cell cycle arrest and epithelial to mesenchymal transition was enriched during enEVT differentiation (Fig.4d). Interestingly, *PRDM6* (PR/SET domain 6) , that played important roles in cell cycle regulation in multiple cell types, for instance vascular endothelial cells and smooth muscle cells based on previous studies ^35, 36^, was highly expressed in enEVT, and the putative target genes involved in cell growth that were suppressed by *PRDM6,* including *HDAC1*, *HDAC3* and *TET1* (Fig. 4e, 4f). Consistent with the above observations, in our study, the immumohistochemical staining showed that PRDM6 coexpression with HLA-G in some cells, while exclusively expressed with HDAC1 in specific cells (Fig. 4g). Based on the above findings, we proposed that *PRDM6* might be a novel regulator in promoting differentiation of enEVT by positive regulation of cell cycle arrest. Collectively, we provided an overview of transcription factors atlas of enEVT subgroups self-renewal, differentiation and invasion, among them, some of TFs involved in cancer cell development regulation, were proposed as putative novel key TFs in promoting EVT subpopulation development. In short, these findings strongly deepening the understanding of the intrinsic regulatory mechanism of EVT subpopulation *in vivo,* although, more work still need to be done for further validation.

**Fig. 4.**
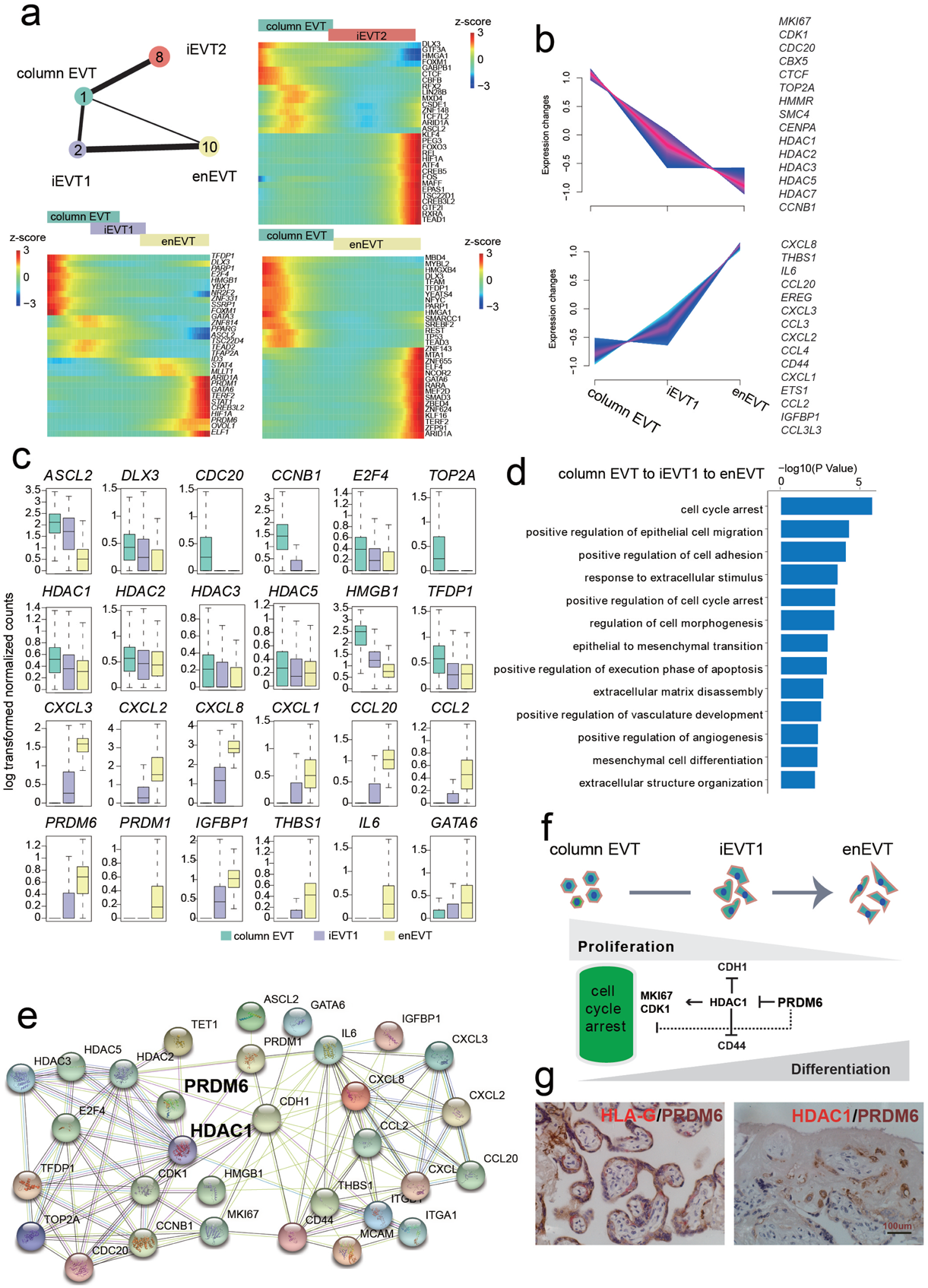
Identification of key transcription factor regulators during extravillous trophoblast cell differentiation. a. Partition-based approximate graph abstraction (PAGA) analysis of EVT subpopulations, including column trophoblast cell (column EVT), interstitial extravillous trophoblast cells 1/2 (iEVT1/2), and endovascular extravillous trophoblast cells (enEVT). Lines show connections; line thickness corresponds to the level of connectivity (low (thin) to high (thick) PAGA connectivity). Heatmap showing min-max normalized expression of statistically significant (P<0.001), dynamically variable transcription factors (TFs) from pseudotime analysis for EVT trajectories. b. The expression pattern of selected DEGs of column EVT, iEVT1 and enEVT. c. Boxplot visualization of log-transformed, normalized expression of selected TFs in EVT subgroups. d. Selected GO terms of TFs differentially expressed in column EVT, iEVT1 and enEVT, respectively. e. Regulatory network of selected TFs differentially expressed in column EVT, iEVT1 and enEVT. f. Model of regulation loops of column EVT differentiation into enEVT. g. Immunostaining of HLA-G, PRDM6 and HDAC1 in Mat_S of human full-term placenta. Scale bar represents 100 μm.

### The transcriptional profiling reveals dysregulation of EVT subgroup in PE

Previous studies showed that abnormal cell type composition and trophoblast differentiation potentially leaded to placental dysfunction and pregnancy complications^37, 38^ . However, the cellular organization of human full-term placenta during both normal and PE development remains largely unknown. In the present study, we combined single cell transcriptome data of both normal placenta from in this study and pregnancy-matched PE placenta from published data^22^ (Supplementary Fig. 5a, 5b, 5c). As expected, genes associated with pregnancy complication from OMIM (Online Mendelian Inheritance in Man) database were differentially expressed in specific cell types between PE and normal groups (Fig. 5a). For instance, *PLAC8*, *PAPPA2*, *FLT1*, *MMP11*, *TAC3*, and *NOS2* were highly expressed in EVT groups; *MMP1*, *EDN1*, *ANGPT2*, *ADAMTS13*, *KLF2*, *NOTCH1*, *LEPR*, *NOS3*, *JAG2*, *SCNN1B* were expressed in VEC cell types (Fig. 5a, 5b). Further, we constructed the regulatory network of pregnancy PE associated genes described above and found that genes such as *FLT1*, *ITGA1*, *EDN1*, *ITGA6*, *ITGB, etc*. were located in core positions (Supplementary Fig. 5d). The above results indicated PE associated genes expression showing pattern specificity and cell type diversity, and suggested that PE is a complicated pregnancy-specific syndrome involving in various cell types and pathways.

**Fig. 5.**
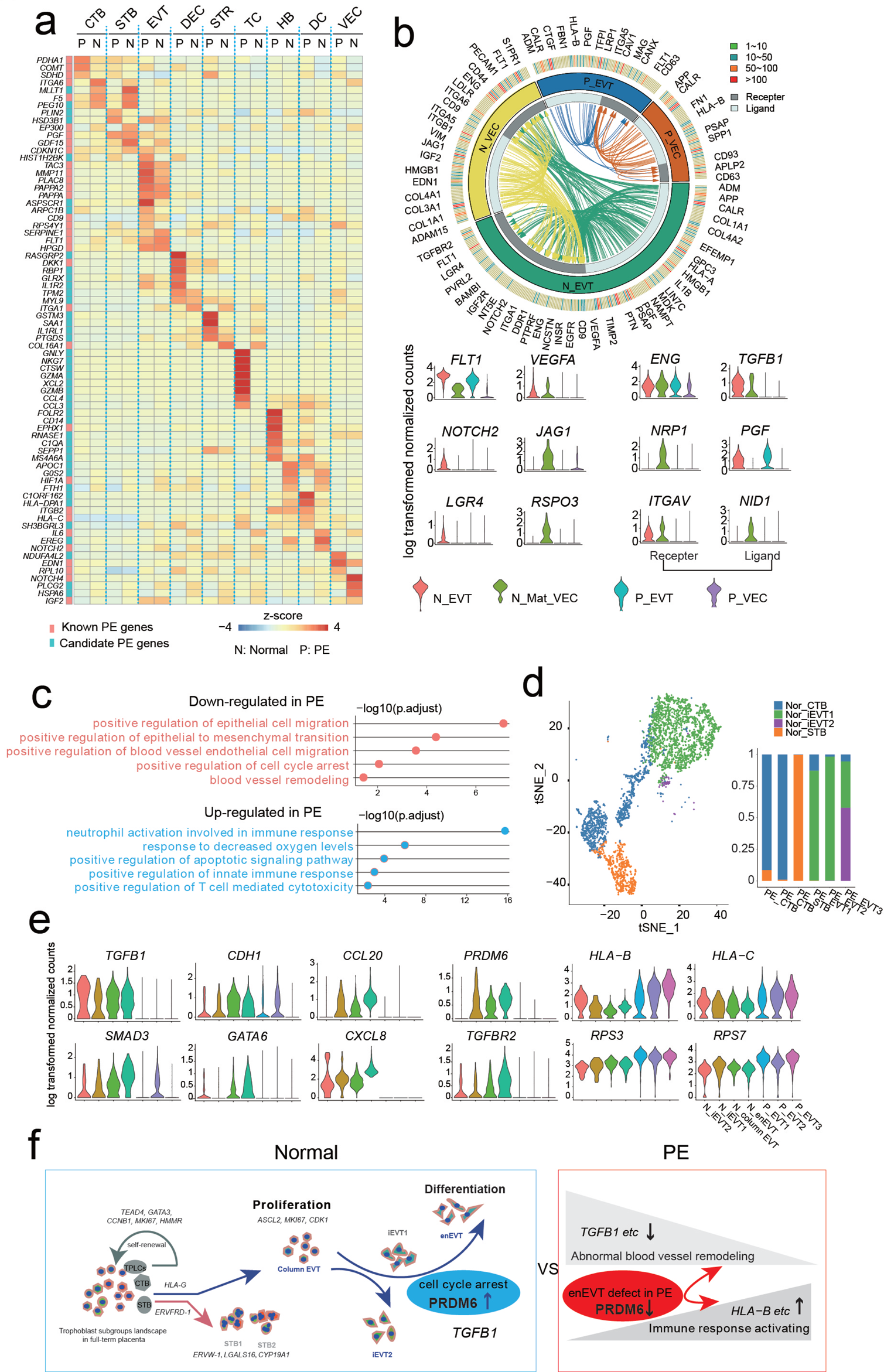
The transcriptional profiling reveals dysregulation of EVT subgroups in PE. a. Heatmap showing the expression level of pregnancy disorder-associated genes downloaded from OMIM website in specific cell types of human normal and PE placenta. b. The ligand-receptor interaction between EVT and VEC in normal and PE samples; genes expressed in more than 40% of cells for specific subtype were selected. Each arrow represents the paired ligand-receptor, and ligands with the same arrow color =belong to the common cell type; violin plots show the selected ligand-receptor pairs for EVT and VEC differentially expressed in normal and PE sample. c. GO term enrichment analysis of genes down-regulated (upper panel) and up-regulated (lower panel) in PE compared to normal placenta. d. The t-SNE plot and column chart showing the consistency of trophoblast subtypes in PE and in normal placenta. e. Boxplot showing the expression level of genes associated with EVT proliferation and differentiation in EVT subgroups between normal and PE samples. f. Proposed schematic of trophoblast subtypes, their self-renewal and differentiation regulated by indicated genes and transcription factors in human normal and PE placenta.

As the EVT populations play crucial role in remodeling VEC to provide ample blood supply to the growing fetus. To further investigate the regulation and communication between fetal EVT and maternal VEC cells, we presented the interaction network of ligand-receptor complexes, which played important roles in vascular remodeling process in both PE and normal placenta, respectively (Fig. 5b). Surprisedly, the ligand-receptor numbers were significantly decreased in PE, e.g., *FLT1-VEGFA, ENG-TGFB1, NRP1-PGF*, *ITGAV-NID1*, and *NOTCH2-JAG1,* belong mainly to terms like “epithelial to mesenchymal transition”. These results above strongly suggested that the EVT development dysfunction and molecules involved in blood vessel remodeling was down-regulated in EVT or VEC of PE placenta.

To systematically dissect the development and cell interaction of EVT in PE. First, the GO analysis for the genes down-regulated in EVT of PE compared to normal sample show terms about “blood vessel remodeling”, “positive regulation of epithelial to mesenchymal transition”, and “positive regulation of cell cycle arrest”; while “neutrophil activation involved in immune response”, “positive regulation of apoptotic signaling pathway” and “positive regulation of T cell mediated cytotoxicity” for the up-regulated genes in EVT of PE (Fig. 5c). Then, the EVT subgroups of PE generally belonged to iEVT1 and iEVT2 subgroups of normal placenta based on the transcriptome mapping analysis using the expression gene matrix of each EVT subgroup (Fig 5d; Supplementary Fig. 5a). And the expression of enEVT differentiation and invasion-related genes and ligand-receptors, such as *ASCL2*, *DIO2*, *ITGA1*, *ITGA5*, *TGAV*, *ITGB1*, *PRDM6* and *CD44* were significantly decreased in EVT subgroups of PE (Fig. 5e; Supplementary Fig. 5e, 5f). In additional, *PRDM6*, as a novel marker gene, was highly expressed in enEVT subgroup in normal condition but significantly reduced in PE (Fig. 5e, 5f). Together with previous report that deficient *PRDM6* was associated with vascular system disease^35^, here, we proposed that the functional dysregulation of *PRDM6*, together with other genes related to EVT differentiation and invasion, may result in placental disorder. In short, these results above suggested that abnormal EVT subgroup composition and defect of invasion or differentiation could be the underlying causes of PE.

## Discussion

Anatomically, the human placenta is a complex and heterogeneous organ consisting of multiple different cell types that carry out varied functions. In the presented work, we firstly generated a comprehensive single-cell transcriptome profiling of the human full-term placenta. Using unsupervised clustering, we identified the trophoblast cell subtypes and non-trophoblast cell types with indication of their fetal or maternal origin and spatial location. In line with previous studies, Mat_S contained mostly maternal cells, e.g., LEC and DEC; while fetal cells such as CTB, EVT, and STB dominate the Mid_S and FS^39^. In additional, IMM mixed with fetal and maternal origin distributed in each section also observed in our study, which was consistent with previous study^40^. Interestingly, proportionate STR, PV, and VEC originated from both fetal and maternal compartments mainly occupied Mid_S. Currently, the interaction between fetal-origin and maternal-origin cells during human placentation and functional maturation is poorly understood. Previous studies showed CTB and STR as the core cell populations presented dynamic molecular feature changing during placental mature progress. In our work, we observed that CTB displayed spatial variation by both molecular expression pattern and function terms from the fetal side to the maternal side in the fetomaternal interface, that strongly suggest that microenvironment of different location of placenta contributed CTB subpopulations cell states or behaviors, and this phenomenon may also can be observed in other tissue and organ^41^.

Moreover, Trajectory analysis revealing that a subpopulation of TPLCs existed in the full-term placenta and mainly distributed in Mid_S, with high expression of cell surface maker *HMMR* and unique molecular features compared to the TPCs derived from first-trimester placenta, which is worth of further investigation. Although, evidence showed human trophoblast progenitor cells probably exist in the full-term placenta and express angiogenic factors^42^. Currently, researchers are not successfully to isolated trophoblast stem/ progenitor cell from full-term placenta that mainly maybe due to unsuitable culture medium in vitro^12^. However, using TPLCs in full-term placenta as an ideal disease model for future research still needs further verification.

Previous studies showed that STR from fetal- and maternal-origin of placenta possess greats differences in biological behaviors, which potential implications for their applications in regenerative medicine^43^. However, insights on the molecular heterogeneity of STR populations within the maternal-fetal interface is still missing. Through DEGs and GO enrichment analysis, we found cells from fetal origins showed greater proliferative ability in Mat_S compared to cells from FS or Mid_S while maintaining the molecular characteristics of the stromal cells, which might be a good resource for mesenchymal stem cell expansion-based cell therapy. And our observes are consisted with previous study showed STR cells are heterogeneous population which caused by growth niche or cell fate decision mechanism.

Normal placental function is dependent on appropriate growth and development of specific cell subsets, which are heterogeneous, dynamic, and are determined by the precise regulation of gene expression. Additionally, through the extravillous cytotrophoblast (EVT) subsets differentiation trajectory and regulation network analysis, we highlighted the putative key transcription factor *PRDM6* that promoted the differentiation of endovascular extravillous trophoblast cells (enEVT). Previous studies showed *PRDM6* played important roles in cell cycle regulation and inhibited vascular endothelial cells proliferation by targeting *HDAC1* gene^35, 36^. Mutations of *PRDM6* are associated with many syndromes due to the abnormal regulation of cell proliferation and apoptosis, such as nonsyndromic patent ductus arteriosus^44^. Combine with the immumohistochemical staining analiysis, we proposed that *PRDM6* might be a novel regulator in promoting differentiation of enEVT by positive regulation of cell cycle arrest.

Consequently, alterations to placental gene expression are thought to be a major cause of pregnancy pathologies. Combined with previously published single-cell transcriptome data of PE, we also highlighted the abnormal EVT subgroup components (enEVT absence) and suggested that the defect of epithelial to mesenchymal transition related ligands and receptors could be the underlying causes of PE. Moreover, we inferred down-regulation of PRDM6 may lead to an abnormal enEVT differentiation process and highly related to PE. However, the reason why column EVT and enEVT are defective in maternal-fetal interface of PE still need to explore. We hope that the trophoblast differentiation cell model *in vitro* combined with single cell omics technology would provide more clues.

Previous studies showed large number of cancer cell features can be recapitulated by development of the placenta^45^ . Among the properties shared by trophoblast cells and cancer cells is the ability to invade healthy tissues, to remodel vessels and to form a niche to regulate immunoreaction^45^. In line with previous study, a large number of cancer cell related TFs, e.g., *SMARCC1*, *GTF3A*, *MYBL2*, *SUB1*, and *NCOR1* etc., contributed to maintaining cancer cell proliferation; whereas TFs like *CREB3L2*, *CEBPB*, *RUNX1* etc., play important roles in EVT differentiation were enriched in specific EVT subpopulations in the present study^46–49 35, 36^. In contrast to cancer-invading cells, EVT cells are eliminated at the end of pregnancy in the maternal tissue^50^. Many of the mechanisms leading to the phenotype of cancer cell are still poorly understood^45^. The study of EVT cells might be useful to understand how cancer cells develop their invasive potential in future study.

In conclusion, we provided a comprehensive understanding of the molecular and cellular map of the maternal-fetal interface of full-term placenta through single cell transcriptome profiling. We found TPLCs existed in full-term placenta with inferred pools of cell surface markers, which is worth of further investigation. Moreover, we compared the transcriptomic difference among stromal cells derived from placenta (including maternal-origins, fetal-origins, different spatial locations), and found that stromal cells from fetal origins in Mat_S showed greater proliferative ability while maintaining the molecular characteristics of the stromal cells, which might be a good resource for mesenchymal stem cell expansion-based cell therapy. Furthermore, combined with previously published single-cell transcriptome data of PE, we inferred down-regulation of *PRDM6* may lead to an abnormal enEVT differentiation process and highly related to PE. Together, this study offers important resources for better understanding of human placenta, stem cells based regenerative medicine as well as PE, and provides new insights on the study of tissue heterogeneity, the clinical prevention and control of PE as well as the fetal-maternal interface.

## Methods

### Ethics statement

The study was approved by the Institutional Review Board on Bioethics and Biosafety of BGI (Permit No. BGI-IRB 19145), and the Shenzhen Second People’s Hospital (Permit No. KS20191031002). The participants signed informed consents and voluntarily donated the samples in this study. Immediately after delivery (between 38-40 weeks of gestation), the intact human placenta tissue samples were collected for further use.

### Collection of human placenta samples

All human full-term placenta tissues were obtained from normal pregnancies after delivery^1^, and samples were transported from hospital to BGI-Shenzhen in an ice box within eight hours. The three parts of whole placenta, including FS, Mid_S, and Mat_S were mechanically separated. Each section then underwent serial collagenase IV(Sigma) and trypsin (Invitrogen) digests, respectively, as previously described with some modifications^20^. Next, single cell suspensions were centrifuged and resuspended in 5 mL of red blood cell lysis buffer (Invitrogen) for 5 min, then the cell suspensions were filtered through a 100 µm cell filter (Corning) and washed twice with phosphate-buffered saline (PBS) (Sigma). After single cell suspension preparation, trypan blue (Invitrogen) staining was used to assess cell viability and cell samples with viability over 90% were used for the following single cell RNA seq experiments.

### Single-cell RNA library preparation and sequencing

Single cells resuspended in PBS with 0.04% bovine serum albumin (BSA) (Sigma) were processed through the Chromium Single Cell 3’ Reagent Kit (10X Genomics) according to the manufacturer’s protocol. Briefly, a total of 10,000 cells per sample were mixed with RT-PCR reagents, and loaded onto each channel with Gel Beads. An average of about 6,000 cells could be recovered for each channel. Cells were then partitioned into Gel Beads in Emulsion in the GemCode instrument, where cell lysis and barcoded reverse transcription of RNA occurred. cDNA molecules were then pooled for amplification and the following library construction, including shearing, adaptor ligation, and sample index attachment. Libraries were sequenced on MGI-seq platform.

### Single-cell transcriptome data preprocessing

Droplet-based single-cell sequencing data were aligned to human genome GRCh38, and barcode and UMI were counted using CellRanger software (Version 2.0.0, 10x Genomics)^51^. Genes that were expressed in less than 0.1% of total cells were removed. Cells with detected gene number of less than 800 or expressed mitochondrial genes of more than 10% were filtered. Moreover, for each library, outliers were detected based on gene number using R function boxplot.stats, and were considered as potential doublets to be removed for downstream analysis.

### Inferring maternal or fetal origin of single cells

We obtained the transcriptome data of three fetal umbilical cord tissues and the whole genome sequencing data of one maternal peripheral blood sample from sample individual c. To get the fetal and maternal-specific SNP arrays, the high-quality sequencing reads were aligned to the human genome GRCh38 using BWA-MEM (Version1.0). Sorting, duplicate marking and single nucleotide variants (SNVs) calling were processed using GATK (Version3.8)^52^. The filter parameters were as follows: QD < 2.0 || MQ < 40.0 || FS > 200.0 || SOR > 10.0 || MQRankSum < -12.5 || ReadPosRankSum < -8.0.

Variants were identified from each cell using the Genome Analysis Toolkit. Briefly, duplicated reads were marked with Picard (Version2.9.2). Next, the recommended SplitNCigarReads was also performed by GATK (Version4.0.5.1). Then BaseRecalibrator and ApplyBQSR algorithms were used to detect systematic errors. At the variant calling step, the HaplotypeCaller algorithm was used to call variants and SelectVariants algorithm was used to select SNP sites. Besides, VariantFiltration algorithm was used to filter the SNP sites with the follow algorithms: --filter-expression “DP < 6|| QD < 2.0 || MQ < 40.0 || FS > 60.0 || SOR > 3.0 || MQRankSum < -12.5 || ReadPosRankSum < - 8.0” --filter-name “Filter” -window 35 -cluster 3.

The fetal and maternal origin of each cell was inferred by our discrimination function (Each section was processed individually). In brief, because only individual c has the corresponding mother blood whole genome sequence variants, this sample was used as our training sample, for which each cell’s fetal or maternal origin was determined by demuxlet^53^ using Cell Ranger-aligned BAM file from FS, Mid_S and Mat_S and WGS VCF file. And then, a fetal SNP dataset reference was built based on the corresponding umbilical single cell RNA sequencing data for each section from our previous study^54^. Using the difference ratio between a single cell SNP and the corresponding fetal SNP dataset reference, we calculated the Ratio of Mahalanobis distance of fetal cells and maternal cells usingthe following formulas:

Ratio = mahalanobis (TstX, mu2, S2)/mahalanobis (TstX, mu1, S1),

While:

mu1 = colMeans(Fet.percent.matrix);

S1 = var(Fet.percent.matrix);

mu2 = colMeans(Mat.percent.matrix);

S2 = var(Mat.percent.matrix);

TstX: The difference ratio between a single cell SNP and the corresponding fetal SNP dataset reference as input Matrix TstX, which included Fet.percent.matrix and Mat.percent.matrix.

Fet.percent.matrix: The difference ratio of individual c’s fetal cell between a single cell SNP and the corresponding fetal SNP dataset reference matrix;

Mat.percent.matrix: The difference ratio of individual c’s maternal cell between a single cell SNP and the corresponding fetal SNP dataset reference matrix;

Then, according to the demuxlet results, the sensitivity, specificity, and accuracy were calculated in different ratio. Finally, the optimal discriminiant ratio was selected based on the sensitivity, specificity, and accuracy for each section. If a cell’s Ratio < fetal discriminiant ratio, the cell was inferred as fetal cell; if a cell’s Ratio > maternal discriminiant ratio, the cell was inferred as maternal cell; otherwise, it was defined as unknow in origin.

### Cell clustering and identification of differentially expressed genes

The standard Seurat v3^55^ integration workflow was used to integrate multiple datasets from each sample to correct batch effects between sample identities. Cell clusters were identified by a shared nearest neighbor (SNN) modularity optimization-based clustering algorithm used in “FindClusters” function in Seurat (Version 3.1.0). Differentially expressed genes were found based on Wilcoxon Rank Sum test using default parameters in “FindAllMarkers” function. The significantly differentially expressed genes were selected with adjusted P value < 0.05 and fold change > 0.25.

### Constructing trajectory

Constructing trajectory and ordering single cells were performed with monocle 2 (Version 2.10.1) using the default parameters^56^. The top 2000 highly variable genes found by Seurat were used. The relationship between each EVT subgroup was inferred by partition-based approximate graph abstraction (PAGA) (Paga in scanpy Python package version 1.2.2).

### GO enrichment analysis

GO enrichment analysis was performed by clusterProfiler R package^57^. The p value was adjusted by BH (Benjamini-Hochberg). GO terms with an adjusted p-value less than 0.05 were considered as significantly enriched.

### Regulatory network construction

Significantly differentially expressed TFs (adjust p value <0.05) between each population were selected and submitted to the STRING database to construct the potential regulatory networks^58^. TFs without any edge were removed from the network.

### Cell–cell communication analysis

The ligand–receptor pairs were obtained from work of Ramilowski et al^59^. A ligand or receptor transcript was selected for a given cell type if it was expressed in more than 40% cells in that cell type. The gene pairs possibly interact on the same cell type were not presented. The interactions were visualized by R package Circlize^60^.

### Integrative analysis of published placenta single cell transcriptome data

The previously reports single cell transcriptome data for first-trimester placentas and the preeclamptic placentas papers published in Nature^20^ and PNAS^22^, were integrated with our data for different analyses. “IntegrateData” function in Seurat V3 were used to remove batch effect.

### Immunohistochemistry

Histologic sections of the normal human full-term placenta were rinsed with xylenes two-three times and rehydrated before labeling. Samples were labeled for 1 h with the primary antibody against MKI67 (1:800 Abcam), KRT8 (1:100 Abcam), CDK1(1:200 Abcam), TPX2 (1:100 Abcam), TEAD4(1:200 Abcam), CCNB1(1:100 Abcam), HLA-G (1:200 Abcam), HDAC1(1:100 Abcam) and PRDM6 (1:200 Abcam) and for 30 min with the secondary antibody goat anti-mouse (1: 500, Abcam) or goat anti-rabbit (1:500 Abcam) as appropriate. Finally, samples were counterstained with hematoxylin to reveal cell nuclei for 1 min. Images were taken by the Olympus IX71 microscope.

## Acknowledgements

We sincerely thank the support provided by China National Gene Bank. This study was supported by Science, Technology and Innovation Commission of Shenzhen Municipality Grant (number JCYJ20180507183628543).

## Data availability

All of the raw data have been deposited into CNSA (CNGB Nucleotide Sequence Archive) of CNGBdb with accession number CNP0000878 (https://db.cngb.org/cnsa/ ).

## Author contributions

Z.S. and W.K. conceived and designed the project. Q.W., J.L. performed the experiments and data analysis, and wrote the manuscript. Q.D., K.W., Y.X., S.W., Y.A., and X.D. prepared figures. G.D., Q.C., Z.L., W.Z., and T.Z. contributed to sample collection and provided suggestions on data analysis. Y.H., D.Y., H.Y. supervised the project. All authors read and approved the final manuscript.

## Figure Legends

**Supplementary Fig 1. Information about the samples and the single-cell datasets quality.**

a. Detailed information of human full-term placenta samples and single cell sequencing data.

b. The density graphic showing the distribution of detected gene number(left), unique feature counts (middle), and the percentage of mitochondrial counts (right)

c. Boxplot showing the expression pattern of canonical marker genes in each cell type.

d. Barplot showing the proportion of each sample in each cell cluster.

e. Table showing the sensitivity, accuracy, and specificity of discrimination

function to infer the origin of fetal or maternal cells in full-term placenta.

**Supplementary Fig 2. Molecular features analysis of STR with specific origin and spatial location.**

a. Selected GO terms identified by top 1000 highly expressed genes in each section of fetal origin (Left) and maternal origin (Right) STR cell. (Highly expressed genes with expressed cell number > 20% and gene coefficient of variability (CV) <1 was used in each section).

b. Boxplot showing the differentially expressed genes of STR cells in each section. Two-sided Wilcoxon rank sum test were calculated, **** p <0.0001.

c. Barplot showing the proportion of STR cells with determined origin and undetermined origin in each section.

d. Heatmap showing the expression pattern of genes related to cytokines and hormones in STR cells from different origin in each section.

e. Boxplot showing the different gene expression between STR cells from different origin in Mat_S. Two-sided Wilcoxon rank sum test were calculated, * p < 0.05, ** p < 0.01, *** p < 0.001, **** p <0.0001.

**Supplementary Fig 3. Integrated data analysis of trophoblast cell from human full-term placenta and downloaded first-trimester placenta.**

a. t-SNE visualization of integrated data for full-term placenta single cell transcriptome data with that from published first-trimester placenta. On the right, barplot shows the proportion of full-term placenta cell and first-trimester placenta cell in each cluster.

b. Violin plot showing the expression of canonical marker genes for the defined cell types. Clusters annotated with the same cell type are shown together. (The clusters in Supplementary Fig 3a that each cell type includes are: CTB: 3, 16, 19, 21; EVT: 8, 11; STB: 29; STR: 7, 9, 13, 32; DEC: 4, 14, 18, 24, 27; PV: 6, 10, 12, 33; VEC: 20; LEC: 23; Dendritic cell, DC: 2, 5, 25; Hofbauer cell, HB: 17, 26, 28; T cell, TC: 1, 15; Natural killer cell, NK: 0, 22; Endometrial Epithelial Cell, EEC: 31)

c. t-SNE Plot showing the expression of canonical marker genes for the defined cell types of re-clustered trophoblast cells shown in Fig. 3a.

d. Location of each trophoblast subgroup of Fig. 3a on the trophoblast cell differentiation trajectory.

**Supplementary Fig 4. The features analysis in each EVT subgroup.**

a. Boxplot showing the expression level of specific genes for each EVT subgroup.

b. Selected GO terms identified by differentially expressed genes for each EVT subgroup.

**Supplementary Fig 5. Comparison of the differentially key features of trophoblast cell subgroups between human normal and PE placenta.**

a. t-SNE visualization of single cell RNA transcriptome data of two selected preeclampsia(PE) placenta in reference 22, colors indicate different cell types or subtypes.

b. t-SNE plot showing the relative expression level of canonical marker genes for the defined cell types.

c. Barplot showing the proportion of each sample in each cellular subgroup.

d. Regulatory network of pregnancy-associated and candidate disease genes from Fig. 5a.

e. Boxplot showing the relative expression levels of genes associated with trophoblast proliferation and differentiation in EVT subgroups between normal and PE sample.

f. Violin plot showing the relative expression levels for selected ligand-receptor pairs in EVT and VEC of normal and PE samples.

